# Mechanical Memory Primes Cells for Confined Migration

**DOI:** 10.1101/2025.09.07.674701

**Authors:** Jia Wen Nicole Lee, Yixuan Li, Xu Gao, Avery Rui Sun, Jin Zhu, Jennifer L. Young, Andrew W. Holle

## Abstract

When migratory cells move from one stiffness niche to another *in vivo*, they are exposed to highly confined spaces imposed by dense extracellular matrix (ECM) networks and inter-tissue boundaries. Cells that originate from one niche possess distinct mechanosensitive adaptations that influence their response to their new niche, a concept known as mechanical memory. However, the mechanisms by which this memory is acquired, and the degree to which it influences migratory potential and decision-making processes in confinement remain poorly understood. Here, we combine stiffness priming using polyacrylamide hydrogels with a confinement platform to screen mechanical memory across healthy and transformed cells. Using a dose-and- passage approach, we find that in stiffness-sensitive cells primed on soft substrates navigate confinement more efficiently. Bulk RNA sequencing identifies NFATC2 as a transcription factor that mediates mechanical memory by reprogramming gene expression in stiffness-sensitive cells. siRNA-induced knockdown of NFATC2 in memory-sensitive cells confirmed its necessity for mechanical memory acquisition and subsequent confined migration enhancement. Interestingly, highly invasive cancer cells exhibit minimal sensitivity to prior mechanical priming, suggesting differential adaptation strategies. These findings reveal mechanical memory as a cell-intrinsic property shaped by past mechanical environments and highlight potential implications for controlling migration in wound repair, fibrosis, and disease progression.

## Introduction

The physical microenvironment *in vivo* poses diverse mechanical challenges to cells, shaping their form and function in processes spanning embryonic development, stem cell homing, wound healing, and immune responses ^1–4^. Mechanical confinement is a common and defining feature in many of these settings, where cells must navigate dense extracellular matrices (ECM), narrow interstitial spaces, and vascular barriers ^5,6^. These spatially restricted environments expose cells to dynamic mechanical cues including matrix stiffness, fluid stresses, and compressive forces, all of which can evolve over distinct timescales ^7^. For instance, matrix stiffening, driven by fibrosis and ECM remodeling, develops gradually over months to years, creating an altered mechanical landscape that influences cell behavior ^8,9^. In contrast, rapid mechanical challenges, such as spatial confinement during migration, exert transient forces that demand immediate cellular adaptation ^10^. The interplay of these cues not only dictates acute cell behavior but may also leave a lasting imprint on cellular function via a process known as mechanical memory.

Mechanical memory, an emerging concept in mechanobiology, describes the ability of cells to retain adaptations to past mechanical environments and apply them in new contexts ^11–17^. For instance, mesenchymal stem cells have been shown to retain their potency after being cultured on soft hydrogels, a process which is mediated by persistent chromatin remodeling and the upregulation of both Yes-associated protein (YAP) and the pre-osteogenic transcription factor RUNX2 ^18,19^. Similarly, numerous studies have highlighted examples of mechanical memory in cell migration, mostly finding that dosing cells on stiff surfaces leads to increased migratory potential ^20,21^. Prolonged culture on stiff substrates can activate integrin-mediated AKT and focal adhesion kinase (FAK) signaling, which in turn drive cytoskeletal remodeling and reinforce actomyosin contractility ^22,23^. Similarly, YAP has been implicated in sustaining mechanical memory via the maintenance of pro-migratory gene expression even after cells are removed from stiff environments ^24,25^. While these and other studies have identified the role of mechanical memory in specific contexts, it remains unclear how prior stiffness dosing can influence cell behavior when navigating confined spaces.

Cells navigating narrow interstitial spaces and dense ECM *in vivo* experience mechanical compression and confinement ^26,27^ and must therefore undergo specific adaptations, including cytoskeletal reorganization, supramolecular complex translocation ^28^, nuclear deformation, and the mesenchymal-to-amoeboid transition ^29–33^. The nucleus in particular plays an important role in this process as it acts as both a physical barrier and a mechanosensor, with extreme deformation triggering nuclear envelope reorganisation and contractility mechanisms to facilitate migration ^34–36^. While cell migration in response to acute confinement has been widely explored ^37^, it remains unclear whether mechanical memory influences the degree to which cells can reorganise and navigate physiological confinement, despite the fact that the vast majority of cells migrating across tissues within the body are moving from one stiffness niche to another.

Here, we investigate how long-term mechanical conditioning as a function of substrate stiffness alters confined migration in both healthy and transformed cells. Transcriptomic profiling of these memory-retaining cells identified a novel candidate for modulating mechanical memory. By perturbing this signaling pathway, we were able to attenuate mechanical memory-influenced confined migration. These findings advance our understanding of how the mechanical microenvironment influences cell migration in a wide variety of settings and has important implications for a number of clinically relevant processes, including wound healing, cancer metastasis, and tissue engineering.

## Results

### Cell Type-Specific Morphological Responses to Mechanical Dosing Reveal Divergent Mechanical Memory Retention

To investigate how prolonged mechanical dosing primes cells to retain a mechanical memory, we first established a screening system to track morphological adaptations of MDA-MB-231 (metastatic breast cancer), HT1080 (fibrosarcoma), and HFF-1 (fibroblast) cells after dosing on varying stiffnesses. These cells were selected due to their mesenchymal migration mode, their diverse migratory capacity and speed, and their varying levels of metastatic potential ^38^. Cells were cultured on polyacrylamide (PA) hydrogels mimicking soft (∼1 kPa) and stiff (∼34 kPa) ECM microenvironments for three or seven days. We utilized a dose-and-passage approach, in which cells are removed from their dosing substrate via trypsin and transferred to glass, in order to identify the consequences of mechanical memory on both the dynamic process of cell spreading and the equilibrium quantity of spread area (**Fig. 1A**). This dosing regimen allowed us to decouple transient stiffness sensing from persistent mechanical memory by observing whether cells retained stiffness-conditioned phenotypes after removal from their original mechanical niche.

**Figure 1.**
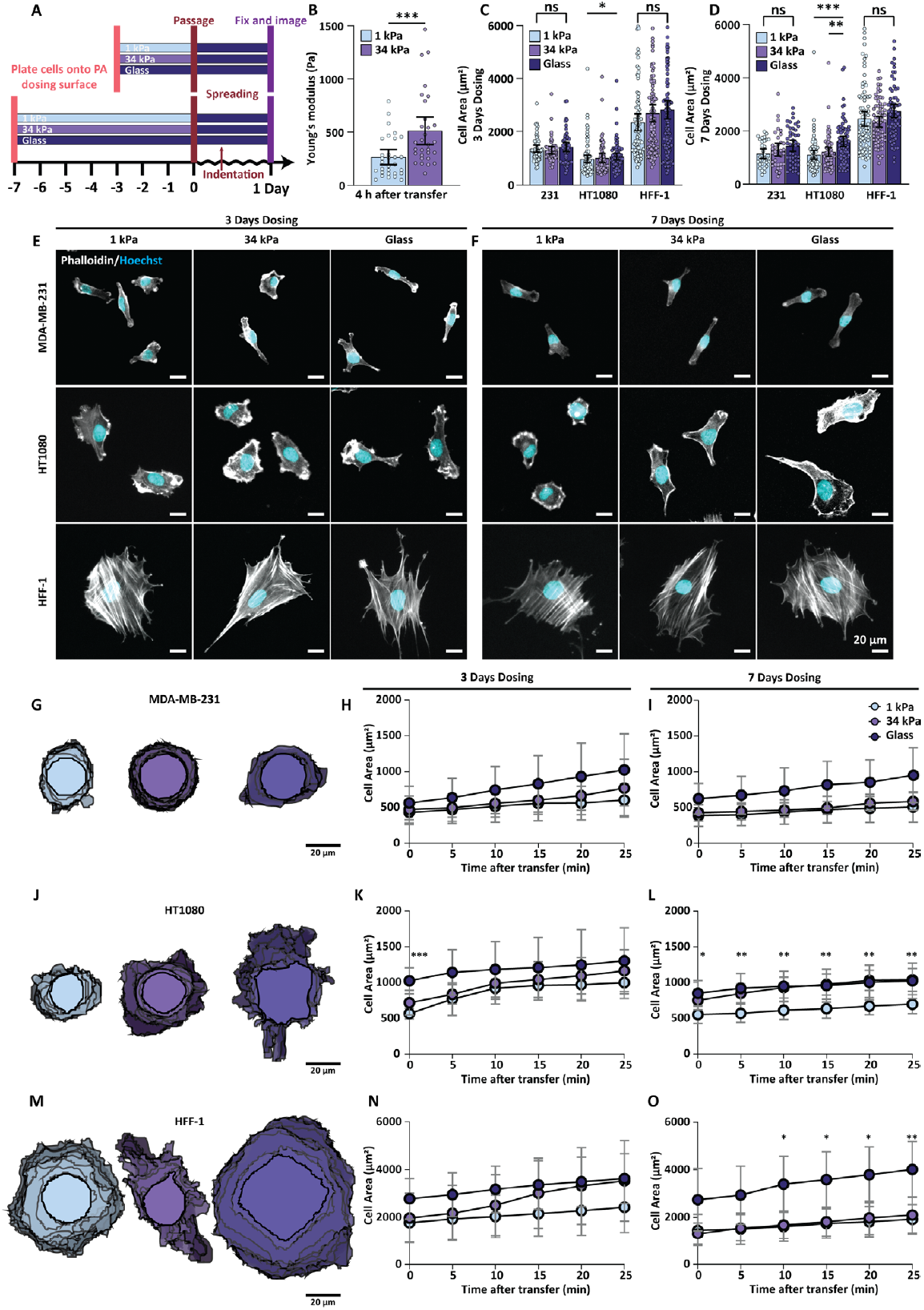
Mechanical Memory-Driven Morphological and Mechanical Responses of MDA-MB-231, HT1080, and HFF-1 Cells to Stiffness Dosing. **(A)** MDA-MB-231, HT1080, and HFF-1 cell lines were cultured on PA hydrogels of different stiffness for 3 or 7 days prior to transfer onto glass surfaces for morphological and stiffness measurements. **(B to D)** Mechanical dosing on varying stiffness for 3 days and 7 days resulted in different cellular stiffness and morphology. Results represented as mean ± 95% C.I., and ^***^ P < 0.001, ^**^ P < 0.01, ^*^ P < 0.05 by Mann-Whitney test or Kruskal-Wallis and Dunn’s post-hoc test. **(E and F)** Example immunofluorescence images of post-dosing cells transferred onto glass coverslips. **(G to O)** Spreading dynamics measured post-mechanical dosing for MDA-MB-231 **(G to I)**, HT1080 **(J to L)**, and HFF-1 cells **(M to O)**, with example overlaid cell boundary maps from 7-day dosed cells. Results represented as mean ± 95% C.I., and ^***^ P < 0.001, ^**^ P < 0.01, ^*^ P < 0.05 by two-way ANOVA and Šídák post-hoc test.

To initially probe mechanical memory, we measured cellular stiffness following hydrogel transfer and found that cells dosed on 1 kPa substrates were significantly softer than those preconditioned on 34 kPa hydrogels (Fig. 1B). Having established this stiffness difference, we next asked whether mechanical memory also manifests as persistent morphological changes. Cells were fixed 24 hours after replating onto glass and systematically assayed for alterations in cell and nuclear morphology (**Fig. 1C-F, Fig. S1A-F**). Following three or seven days of dosing on soft or stiff hydrogels, HFF-1 and MDA-MB-231 cell area did not vary as a function of prior stiffness (**Fig. 1C-D**). In contrast, HT1080 cells preconditioned on soft hydrogels for three days displayed a reduction in cell area of roughly one third compared to those dosed on stiff substrates or glass controls (**Fig. 1C,E**). This effect was even more pronounced after a longer seven day dosing period (**Fig. 1D,F**). Across all cell types and dosing conditions, nuclear aspect ratio (**Fig. S1B,E**) and cell aspect ratio (**Fig. S1A,D**) remained consistent, suggesting that the cytoskeleton is the primary morphological indicator of mechanical memory. Together, this shows that HT1080 fibrosarcoma cells are capable of encoding long-term mechanical memory through sustained alterations in cell spreading.

While cell morphology at a single time point following dose-and-passage can indicate the presence of durable mechanical memory, it does not reveal the dynamics of cell spreading following trypsin-induced cytoskeletal reorganization. To understand this, we observed Lifeact-labeled cells attaching to and spreading on glass after three or seven days of mechanical dosing (**Fig. 1G-O**). Similar to observations made after 24 hours, both HFF-1 cells and MDA-MB-231 cells did not show strong memory-dependent cell spreading profiles (**Fig. 1G-I, 1M-O**). On the other hand, HT1080 cells preconditioned on soft hydrogels spread roughly 20% slower than those on stiff hydrogels (**Fig. 1J-K**), revealing that mechanical memory can play a role in spreading events as early as 25 minutes post-passage. This effect was magnified when HT1080s were dosed for seven days, showing that mechanical memory can accumulate as a function of time (**Fig. 1L**).

### Stiffness-primed mechanical memory drives differential migratory behavior

To assess whether mechanical memory influences persistent motility, we tracked random migration of mechanically preconditioned cells (**Fig. S2A)**. Mirroring their insensitivity to mechanical dosing in prior assays, MDA-MB-231 cells exhibited no differences in migration speed or directionality across dosing conditions or durations, while HT1080 and HFF-1 cells showed robust mechanical memory phenotypes (**Fig. 2A-D, Fig. S2B-E**). Soft-dosed populations migrated over 50 percent faster than the glass negative controls after three days, with this difference increasing in magnitude after seven days of preconditioning (**Fig. 2E-H, S2F-I**). These results parallel earlier morphological and spreading trends, confirming that prolonged mechanical dosing stabilizes memory in fibroblasts and fibrosarcoma cells, while metastatic cells remain indifferent to prior mechanical history.

**Figure 2.**
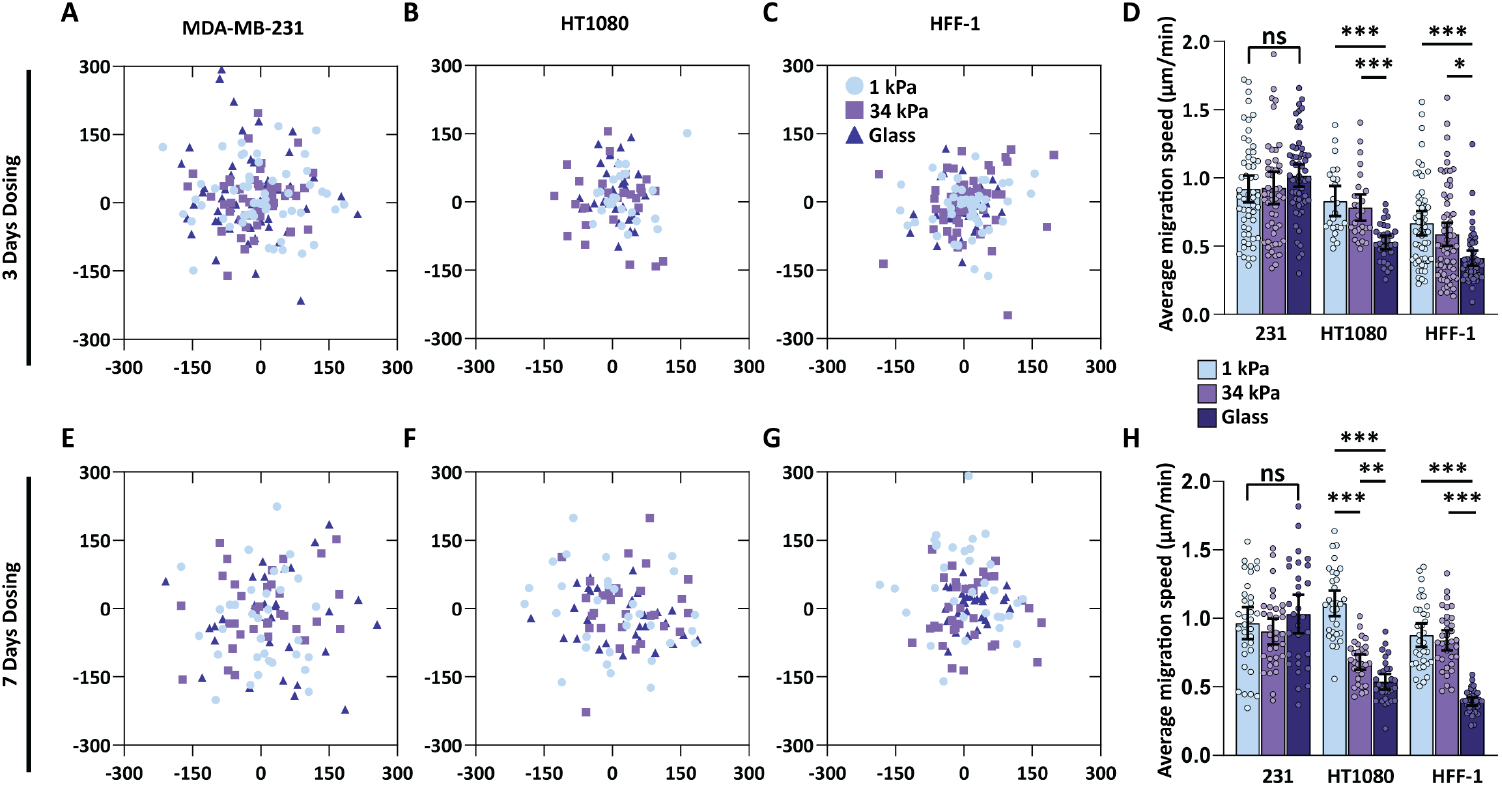
Mechanical Memory-Driven 2D Migration Responses of Cells to Stiffness Dosing. **(A to C, E to G)** Scatter plots mark the 2D migration trajectory end point of MDA-MB-231 **(A and E)**, HT1080 **(B and F)**, and HFF-1 cells **(C and G)** post mechanical dosing. Unit: µm **(D and H)** Average migration speed measured across cell lines with 3-day **(D)** and 7-day **(H)** mechanical dosing. Results represented as mean ± 95% C.I., and ^***^ P < 0.001, ^**^ P < 0.01, ^*^ P < 0.05 by one-way ANOVA and Dunnett’s post-hoc test.

While many studies have shown mechanical memory with morphological measurements and random migration as a readout, confinement is an underrepresented and unique approach to assessing mechanical memory acquisition. Seven day-dosed cells were seeded into PDMS microchannel arrays containing channels of either 3 µm or 10 µm widths, representing a range of physiological levels of confinement observed *in vivo* ^26,27^ (**Fig. 3A-B**). After 24 hours, MDA-MB-231 cells exhibited no difference in migration success as a function of dosing conditions, regardless of channel width (**Fig. 3C, D**), further highlighting their mechanical amnesia. In contrast, preconditioned HT1080 and HFF-1 cells displayed a striking confinement-specific response in which soft-dosed populations were more efficient at permeating narrow 3 µm channels compared to their stiff-dosed counterparts (**Fig. 3C, E-F**). This memory-induced migration enhancement was absent in wider 10 µm channels, suggesting that soft mechanical dosing optimizes cells for entrance and permeation through extreme confinement by facilitating cytoskeletal or nuclear adaptations critical for navigating subnuclear-sized spaces.

**Figure 3.**
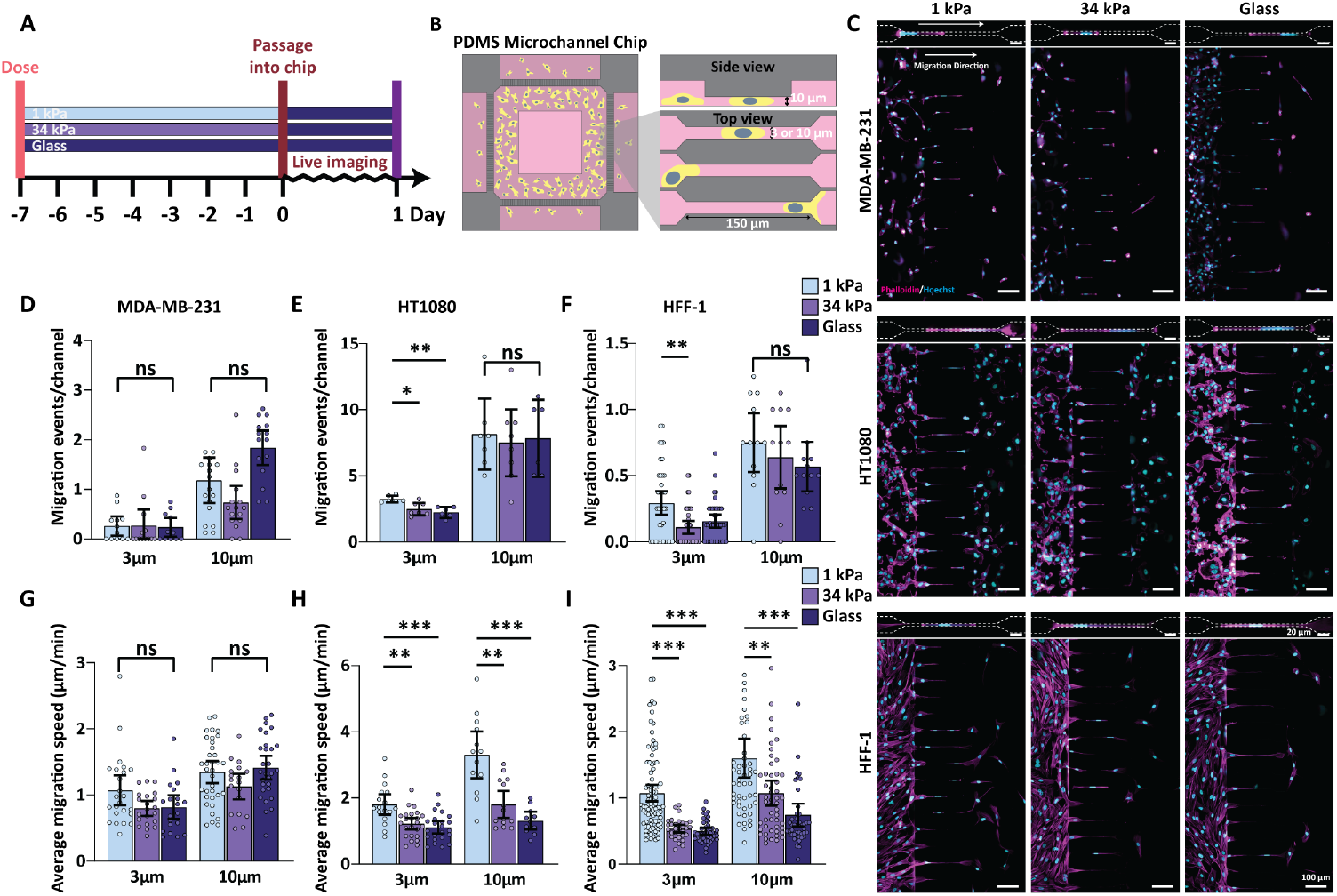
Confined Migration as a Functional Output of Cellular Mechanical Memory. **(A and B)** Cells were introduced into 3 µm-wide and 10 µm-wide microchannels after mechanical dosing for 7 days, and live-imaged over the course of 24 hours. **(C)** Representative immunofluorescence images of migration events through 3 µm-wide confining microchannels 24 hours post-seeding. **(D to I)** Quantification of successful confined migration events per channel over 24 hours for the three mechanically dosed cell lines through both 3 µm-wide and 10 µm-wide channels. Results represented as mean ± 95% C.I., and ^***^ P < 0.001, ^**^ P < 0.01, ^*^ P < 0.05 by one-way ANOVA and Dunnett’s post-hoc test.

We next asked whether mechanical memory influences not only migration success but also the speed of confined movement. While MDA-MB-231 cells again showed no stiffness-dependent differences in migration speed across all channel widths (**Fig. S3A, Fig. 3G**), HT1080 and HFF-1 cells exhibited a striking mechanical memory-dependent effect, with soft-dosed populations migrating over 60% faster than their stiff-dosed counterparts in both 3 µm and 10 µm channels (**Fig. S3A, Fig. 3H-I**). Combined with our earlier finding that soft dosing specifically enhances 3 µm channel permeation, these results imply that mechanical memory can facilitate both adaptation to confinement (enabling entrance and passage through subnuclear spaces) and motility potentiation (increasing cell speed in both confined and unconfined settings). MDA-MB-231 cells’ indifference to mechanical dosing further underscores their defective mechanosensory machinery.

### Bulk RNA sequencing reveals NFATC2 as a common upregulated gene in cells that retain mechanical memory

Transformed and metastatic cells like MDA-MB-231s have been shown to lack essential components of the molecular machinery required for sustained stiffness sensing ^39^, which may explain their inability to retain mechanical memory. To identify conserved regulators of this phenomenon, we focused on transcriptional programs persisting beyond transient cytoskeletal rearrangements. Our dose-and-passage approach ensured that only stable, memory-encoding pathways (e.g., transcriptional/epigenetic changes) were captured, as cytoskeletal adaptations dissipate within hours post-trypsinization ^40^. Consistent with this, nanoindentation revealed that stiffness differences between 1 kPa- and 34 kPa-dosed cells were lost by day 3 (**Fig. S4D-E**). Bulk RNA sequencing was therefore performed after passaging the dosed cells, comparing cells preconditioned on 1 kPa hydrogels against glass-dosed controls to highlight mechanical memory-driven differential expression (**Fig. 4A-B**).

**Figure 4.**
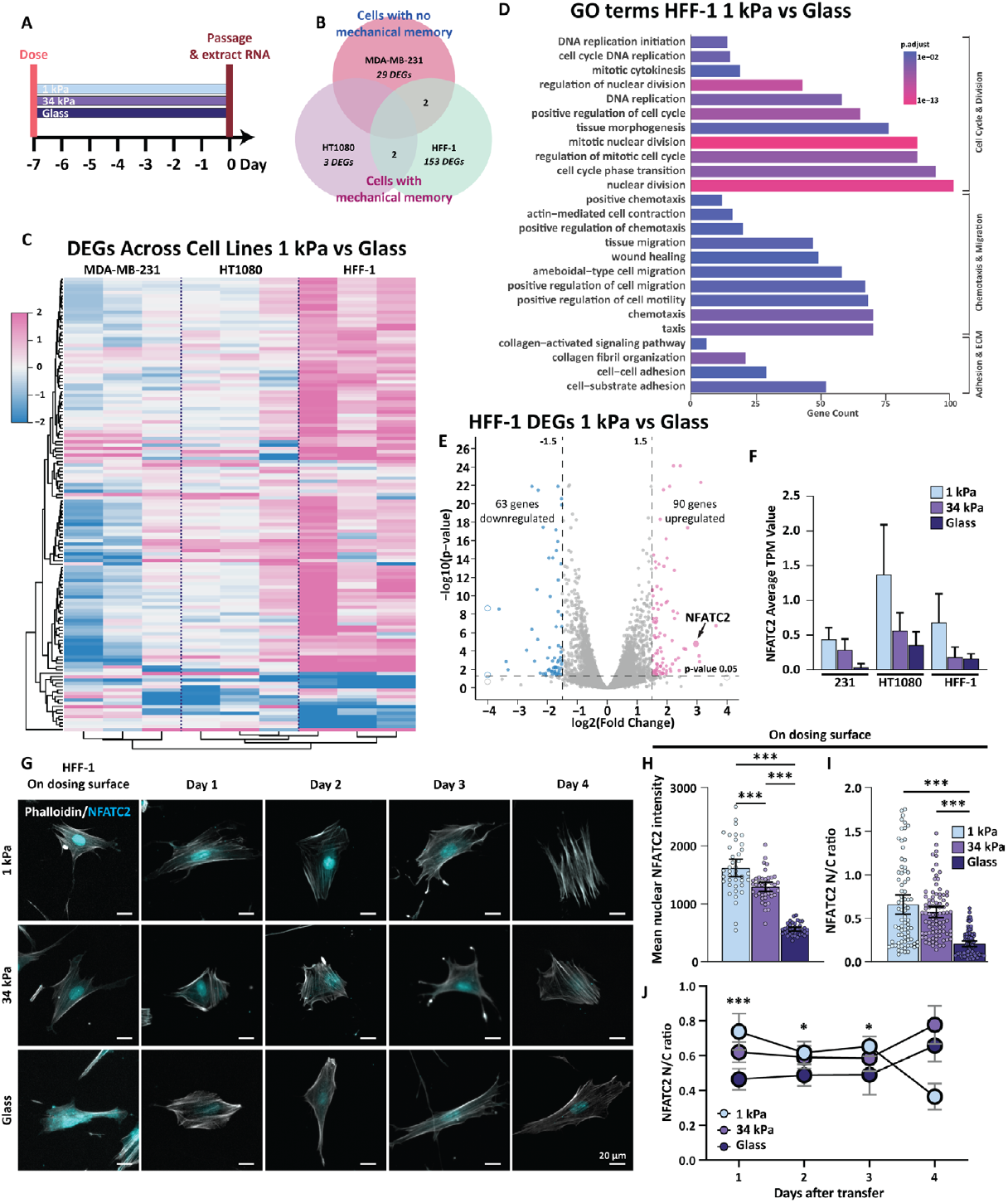
RNA Sequencing Reveals NFATC2 as a Key Regulator of Mechanical Memory. **(A)** Cells dosed for 7 days were collected for RNA extraction and sequencing. **(B)** Venn diagram of the three cell lines showing shared differentially expressed genes (DEGs) from 1 kPa-dosed cells compared to glass. **(C)** Heatmap showing the expression level of DEGs shared across cell lines resulting from 7-day mechanical dosing on 1 kPa hydrogel surface compared to glass. **(D)** Differentially enriched gene ontology (GO) pathways from dosed HFF-1 cells on 1 kPa hydrogel surface compared to glass. **(E)** Volcano plot showing differentially expressed genes between HFF-1 cells dosed on 1kPa hydrogel surface and glass. Points cut off by axis were indicated by open circles. **(F)** NFATC2 gene transcripts per million (TPM) measured from MDA-MB-231, HT1080, and HFF-1 cells with different mechanical dosing. Results represented as mean + SD. **(G)** Example fluorescence images showing NFATC2 protein localization changes in dosed HFF-1 cells over the course of 4 days. **(H and I)** NFATC2 mean nuclear intensity and nuclear-to-cytoplasmic (N/C) ratio in dosed HFF-1 cells. Results represented as mean ± 95% C.I., and ^***^ P < 0.001 by Kruskal-Wallis and Dunn’s post-hoc test. **(J)** NFATC2 N/C ratio changes in dosed HFF-1 cells over the course of 4 days post-dosing. Results represented as mean ± 95% C.I., and ^***^ P < 0.001, ^*^ P < 0.05 by mixed effect analysis.

Pathway analyses highlighted enrichment in pathways related to cell division (e.g., “mitotic nuclear division”), cell migration (e.g., “amoeboidal-type cell migration”), and ECM remodelling (e.g., “collagen fibril organization”) (**Fig 4D, Fig. S4A-B)**. Cross-comparison of differentially-expressed genes (DEGs) between HT1080s and HFF-1s identified two shared genes: the downregulated HISTH2BC gene, a histone variant with uncharacterized mechanosensitive roles, and the upregulated NFATC2 gene, a calcineurin-responsive transcription factor implicated in cell migration and cancer progression ^41^ (**Fig. 4C, E, Fig. S4C**)

NFATC2 emerged as the most differentially expressed gene (log2FC = 1.98 in HT1080s, 2.97 in HFF-1s), with no significant upregulation in MDA-MB-231s (**Fig. 4F**). While NFATC2’s role in mechanotransduction is unknown, it is a plausible mediator of the observed mechanical memory phenotypes. Its stiffness dose-dependent expression, which is absent in metastatic cells, suggests that fibroblasts and fibrosarcoma cells leverage NFATC2 to retain mechanical memory, while metastatic cells bypass such regulation to maintain plasticity in heterogenous environments.

Immunofluorescence staining confirmed the RNAseq findings that NFATC2 is differentially regulated in mechanically-dosed cells. Specifically, we found a striking stiffness-dependent localization pattern of NFATC2 in HFF-1s. On soft hydrogels, NFATC accumulated heavily in the nucleus, where it is likely transcriptionally active, while stiff substrates promoted cytoplasmic retention (**Fig. 4G-I**). After replating onto glass, nuclear localization of NFATC2 in soft-dosed cells gradually diminished over 72 hours (**Fig. 4G,J**). This spatiotemporal regulation suggests that soft mechanical dosing drives NFATC2 nuclear translocation, initiating transcriptional programs that persist post-replating to sustain memory, before the ‘loss’ of mechanical memory occurs 3-4 days after transfer to a new microenvironment.

### Knockdown of NFATC2 in HT1080 and HFF-1 cells results in the attenuation of mechanical memory

To confirm that NFATC2 is a necessary regulator of mechanical memory, we inhibited its expression in HT1080 and HFF-1 cells using siRNA (**Fig. 5A-C**). Even with knockdown levels of roughly 30%, reduction of NFATC2 resulted in a complete loss of the stiffness-dependent migration advantage that had been conveyed onto soft-dosed populations. In both 3 µm and 10 µm microchannels, soft-dosed HT1080 and HFF-1 cells migrated at speeds similar to glass-dosed control cells (**Fig. 5D-G**). These results confirm that NFATC2 is a necessary mediator of mechanical memory, enabling fibroblasts and fibrosarcoma cells to retain stiffness-conditioned cell adaptations.

**Figure 5.**
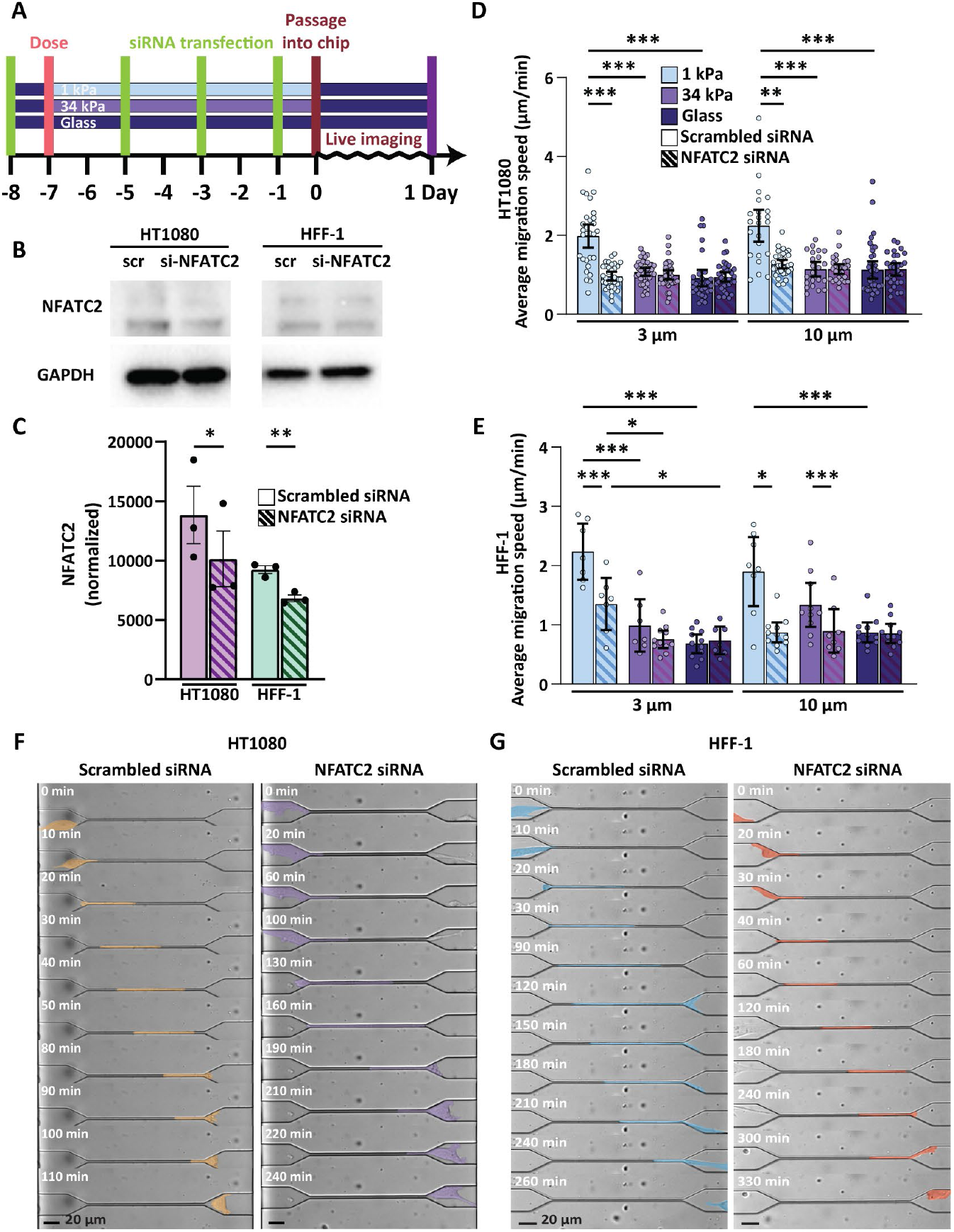
NFATC2 Knockdown Attenuates Mechanical Memory in Cells. **(A)** siRNA knockdown was performed prior to and during 7-day mechanical dosing on hydrogels of different stiffness. **(B and C)** Western blot analysis and quantification of NFATC2 knockdown. Results represented as mean ± SEM, and ^***^ P < 0.001, ^**^ P < 0.01, ^*^ P < 0.05 by paired t-test. **(D and E)** Measured average migration speed of control and NFATC2 knockdown HT1080 **(D)** and HFF-1 **(E)** cells through 3 µm-wide and 10 µm-wide microchannels. Results represented as mean ± 95% C.I., and ^***^P < 0.001, ^**^ P < 0.01, ^*^ P < 0.05 by Kruskal-Wallis and Dunn’s post-hoc test. **(F and G)** Time-lapse images of individual dosed control and NFATC2 knockdown HT1080 **(F)** and HFF-1 **(G)** cells through 3 µm-wide microchannels. Colored stamps indicate cell outlines.

## Discussion

In the body, a wide variety of cells are constantly on the move through various levels of physical confinement, both in routine homeostasis and in highly specific disease contexts. Regardless of whether this movement is inter-tissue or intra-tissue, this confined migration almost by definition brings cells from one mechanical microenvironment to another. While this is most evident in metastasizing cancer cells, which move from their stiff, dysregulated tumor microenvironment into distant tissue, it also occurs in immune cell homing (from lymph nodes to sites of inflammation) and in fibroblast activation (from sites of quiescence to wound healing environments). As this journey necessitates exposure to different extracellular matrix properties both before and after movement, the concept of mechanical memory is deeply entwined with confined migration *in vivo*. The possibility that this mechanical memory may even prime their behavior during and after confined migration is especially appealing, as it offers an additional mechanism through which cell behavior can be understood and ultimately controlled.

Here, we demonstrate that persistent mechanical memory can directly influence the ability of cells to undergo confined migration in a cell type-specific fashion, even in the absence of memory-induced morphological changes. This allows for the use of confined migration as a functional readout of mechanical memory acquisition, one that is more robust than observational readouts like cell morphology.

Taking advantage of this, we used bulk transcriptomics to identify differentially regulated gene expression as a function of stiffness dosing and identified which genes play a key role in the induction of mechanical memory and subsequent memory-specific confined migration. Based on its significant upregulation in memory-possessing cell types, we identified the transcription factor NFATC2 as a key and novel mediator of mechanical memory, diverging from its canonical roles in the epithelial-to-mesenchymal transition, invasion, and survival in cancer cells ^42–45^. NFATC2 is known to shuttle between the nucleus and the cytoplasm based on calcineurin-mediated dephosphorylation (cytoplasmic export) or kinase-driven rephosphorylation (nuclear retention) ^46^, which dynamically modulates its function and could possibly regulate mechanical memory dissipation. In line with this, our RNA sequencing results also revealed differential expressions of several genes involved in calcium signaling, including CAMK1G, TRPC6, GNA14, CALML4, NALCN, and multiple RGS family members. These findings suggest that calcium signaling pathways may act in parallel with or upstream of NFATC2 to encode mechanical memory.

While NFATC2’s function in mechanosignaling is not well known, increased expression levels have been correlated with integrin α6β4 activity in MDA-MB-435 breast cancer cells, leading to increased invasion behavior through transwells assays ^47^. Interestingly, recent studies have shown that YAP acts as a negative regulator of NFATC2 activity in T cells, through sequestration in the cytoplasm via an IQGAP1-mediated complex that prevents NFATC2 nuclear translocation under soft mechanical conditions ^48^. This supports the idea that NFATC2 localization is under mechanical control and may be tightly coupled with cytoskeletal tension and upstream mechanotransducers. These findings position NFATC2 at the intersection of mechanotransduction and calcium signaling, linking soft-induced mechanosensitive cues to a transcriptional response that reinforces or sustains mechanical memory.

This agrees well with our finding that elevated NFATC2 expression in soft-dosed cells resulted in improved migration through microchannels presenting physiological confinement. Critically, our work extends NFATC2’s role beyond classical invasion. Its ability to record prior mechanical experiences via its molecular regulation confers an increased migratory phenotype in confined migration after soft dosing. Recent studies have also demonstrated that Piezo1-mediated calcium influx can be harnessed to drive NFAT-based gene expression for remote-controlled CAR-T activation, further supporting NFATC2 as a promising therapeutic target ^49^. Taken together, this positions NFATC2 not only as a therapeutic target for cancer but as a mediator of context-dependent mechanoadaptations, where a cell’s mechanical history can be leveraged to inform clinical intervention strategies.

It should be noted that mechanical memory operates across a wide spectrum of timescales, shaped by the molecular processes that encode, store, and dissipate it ^13^. As a direct result of mechanosensing, rapid cytoskeletal rearrangements including actomyosin contractility, polymerization, or focal adhesion turnover that adapt cells to mechanical cues occurs at timescales much shorter than other molecular regulations. These changes, while critical for immediate responses, are inherently transient, dissipating once the stimulus is removed and is an unlikely source of long-term mechanical memory storage. In contrast, transcriptional reprogramming lasts hours to days and enables cells to consolidate mechanical signals into sustained functional changes. At the longest timescales, epigenetic modifications lock in mechanical experiences, enabling memory persistence across cell divisions. Positive feedback between these layers of molecular regulations further amplifies and prolongs memory, and the interplay between them can determine whether mechanical memory manifests as a fleeting adaptation or a lasting phenotype. In our dose-and-passage system, cells by definition lose their anchorage to the substrate, which does not occur in situations in which cells remain attached while their mechanical microenvironment changes. This would include disease processes *in vivo* (e.g. tissue fibrosis ^3,50^ or tumor progression ^51–53^ as well as engineering approaches in vitro (e.g. static durotactic gels ^54^, dynamic hydrogel stiffening ^55–59^. However, there are many physiological instances in which cells do detach from their substrate and move into new environments, most notably during cancer metastasis and as a direct result of the adherent-to-suspension transition (AST) ^60^. In addition to this, the dose-and-passage approach ensures that mechanical memory is not a function of the simple rearrangement of the cytoskeleton over time, but rather a more complex integration of mechanosensitive signals that persist even after cytoskeletal rearrangement and reformation.

While these results establish confined migration as a robust functional readout of mechanical memory, the mechanical cues that cells are exposed to during their confining journey also requires new levels of mechanoadaptation. Confinement itself may act as a mechanical imprint, altering cellular function long after the physical constraint is relieved. For instance, transient confinement could epigenetically prime stem cells toward specific lineages^61^ or perpetuate fibrotic activation in fibroblasts, akin to how stiffness memory sustains migratory phenotypes. This bidirectional relationship, in which cells encode past mechanical experiences to navigate future confinement, and confinement then reshapes future cellular behavior, suggests a dynamic reciprocity between mechanical input and output. Future studies will explore whether confinement-induced memory operates through mechanisms like nuclear deformation, persistent cytoskeletal adaptations, or chromatin remodeling, and how these processes intersect with disease states where confinement is a hallmark (e.g., fibrosis, tumor intravasation). This will require the development of new high-throughput systems that can allow for the collection of large populations of cells post-confinement. Ultimately, unraveling confinement’s dual role as both a consequence and catalyst of mechanical memory may redefine how we model cellular adaptation in evolving physiological landscapes.

## Lead contact

Requests for further information and resources should be directed to the lead contact, Andrew W. Holle

## Materials availability

This study did not generate new unique reagents

## Data and code availability

All raw RNA sequencing data have been deposited in the NCBI Sequence Read Archive (SRA) under BioProject accession number [PRJNA1258606]. The data will be made publicly available upon publication.

## Acknowledgments

The authors acknowledge the support of the National Research Foundation of Singapore under an NRF Fellowship to AWH (NRFF13-2021-0114). The authors thank Yeji Chang for fruitful discussions. We are grateful to Gianluca Grenci and Mona Suryana of the Nano and Micro Fabrication Core Facility of the Mechanobiology Institute (MBI) for their assistance with the fabrication of silicon wafers and Paramasivam Kathirvel of the High-Throughput Molecular Genetics Core Facility, MBI for providing input on the knockdown experiments.

## Author contributions

AWH and JWNL conceptualized and designed the study and wrote the manuscript. JWNL, YL, XG, ARS, and JZ performed experiments. JWNL analyzed and visualized the data. JLY and AWH supervised the study.

## Supporting Information

Document S1. Figures S1-S4.

## Materials and methods

### Cell culture

HFF-1, MDA-MB-231, and HT1080 cell lines were obtained from ATCC and were regularly screened for Mycoplasi ma contamination. LifeAct-labelled HFF-1 and HT1080 cell lines were also generated via lentiviral transduction using Lenti-X^™^ Packaging (Takara Bio) with 5 µg/mL polybrene (Sigma-Aldrich) using LifeAct-TdTomato61 lentivirus plasmids (Addgene) while LifeAct-labelled MDA-MB-231s were a gift from Dr. Lanfeng Liang. Fluorescently labeled cells were then sorted using a Sony SH800S Cell Sorter. Cells were maintained in a humidified incubator at 37^°^ C with 5% CO_2_ and were passaged upon 70– 80% confluence. All cell lines were cultured using Dulbecco’s Modified Eagle Medium (DMEM, GIBCO), supplemented with 10% Fetal Bovine Serum (FBS, GIBCO) and 1% Penicillin-Streptomycin (GIBCO).

### Hydrogel Fabrication and Mechanical dosing

Polyacrylamide hydrogels with stiffness values of 1 kPa and 34 kPa were fabricated following a modified protocol based on Tse and Engler ^62^. Briefly, 30 mm coverslips were treated with a UV/ozone cleaner (Bioforce ProCleaner Plus) for 5 minutes, then immersed in a solution containing ethanol, 10% acetic acid, and 3-trimethoxysilylpropylmethacrylate (Sigma Aldrich) for 5 minutes. Coverslips were washed twice in ethanol, air-dried, and prepared for hydrogel adhesion. Chloro-silanized glass slides were prepared by vapor-depositing ∼100 µL of dichlorodimethylsilane (DCDMS, Sigma Aldrich) onto each slide in a fume hood, ensuring full surface coverage. After a 5-minute reaction, excess DCDMS was removed with a Kimwipe, and slides were rinsed with distilled water for 1 minute. Hydrogels were polymerized between methacrylated coverslips and silanized slides using a 200 µL solution of acrylamide and bisacrylamide (Bio-Rad), with Ammonium Persulfate (Bio-Rad) and TEMED (Sigma Aldrich) initiating free radical polymerization. The hydrogels were then washed with PBS before adding 500 µL of 0.2 mg/mL sulfosuccinimidyl-6-(4′-azido-2′-nitrophenylamino)-hexanoate (sulfo-SANPAH, SciHub) solution to coat the gel surface. The gels were exposed to 365-nm UV light (UV-KUB 9, KLOÉ SA) for 3 minutes at 5% power to activate crosslinking. Excess sulfo-SANPAH was removed by washing with PBS, followed by the addition of rat-tail collagen-I (GIBCO) at a concentration of 0.3 mg/mL. Hydrogels were washed with PBS three times before cell culture use. For dosing, cells were seeded at a density of 1,000 cells/cm^2^ for HT1080 and MDA-MB-231 cell lines and 5,000 cells/cm^2^ for HFF-1 cells. Cells were cultured on the hydrogels for 3 or 7 days with the culture medium refreshed after 3 days. At the end of the dosing phase, cells were detached from the hydrogels using 0.05% trypsin (GIBCO) and continuous shaking before subsequent downstream assays.

### Microchannel Chip Fabrication

Wafers containing custom microchannel arrays measuring 150 µm in length, 10 µm in height, and either 3 µm or 10 µm in width were produced by the Microfabrication Core at the Mechanobiology Institute (MBI). Silicon masters were fabricated using a two-step photolithography process. Briefly, wafers were dehydrated at 180 ^°^ C for 15 minutes before spin-coating with a 10 µm SU-8 3010 layer (2900 rpm, 30 s). The wafers were soft-baked (65 ^°^ C for 1 min, 95 ^°^ C for 5 min), exposed to 110 mJ/cm^2^ UV light through a sodalime optical mask (SUSS Microtec MJB4, 365 nm filter), and post-baked (65 ^°^ C for 2 min, 95 ^°^ C for 5 min). After development with SU-8 developer, they were rinsed with isopropyl alcohol, dried with nitrogen gas, and hard-baked at 130 ^°^ C for 5 minutes with cooldown steps at 95 ^°^ C and 6 5^°^ C. A second 100 µm SU-8 3050 layer was applied (1200 rpm, 30 s), soft-baked (65 ^°^ C for 10 min, 95 ^°^ C for 20 min, 65 ^°^ C for 10 min), exposed to 150 mJ/cm^2^ UV light, and post-baked (65 ^°^ C for 2 min, 95 ^°^ C for 5 min, 65 ^°^ C for 2 min). The developed wafers were rinsed, dried, and hard-baked. All SU-8 photoresists and developer were obtained from Kayaku Advanced Materials (MA, USA). Sylgard 184 polydimethylsiloxane (PDMS) (Dow-Corning) was prepared by mixing the polymer base with a curing agent at a 10:1 volumetric ratio, followed by degassing under vacuum for 5 minutes to remove air bubbles. The mixture was then poured over the silicon wafer mold and cured at 80 ^°^ C for 2 hours prior to excision of the negative microchannel structures. Structured PDMS and glass coverslips were plasma treated and attached together to form microchannel chips which were subsequently coated with 0.3 mg/mL rat-tail collagen-1 (GIBCO) dissolved in 0.1% acetic acid overnight at 37 ^°^ C.

### Cell spreading and migration assays

Time-lapse microscopy was performed using the ZEISS Celldiscoverer 7 microscope with a Plan-APOCHROMAT 20×/0.7 NA autocorrective objective. LifeAct-labeled cells were seeded onto collagen-coated glass coverslips after mechanical dosing and fluorescent time-lapse images were acquired every minute immediately after seeding to measure cell spreading events. For cellular migration assays, mechanically-dosed cells were allowed to adhere to the glass coverslip for 1 hour, followed by phase-contrast microscopy images acquired every 10 minutes for 5 hours. Average speed, directionality, and track plots were obtained using the Chemotaxis and Migration Tool V2.0 from Ibidi. For confined migration assays, mechanically-dosed cells were seeded into the entrance reservoir of the PDMS microchannel devices, and migration events were tracked for 24 hours by phase-contrast time-lapse images acquired every 10 minutes.

### Nanoindentation

An Optics11Life Chiaro nanoindenter was used for cell measurements with a spherical probe (k = 0.019 N/m, tip radius = 2.5 mm). Mechanically-dosed cells were plated on glass bottom dishes for 4 hours or 3 days and stained with Hoechst before nanoindentation experiments. A humidified 37 ^o^C chamber built around the nanoindenter was used during measurements. The probe was then mounted on the nanoindenter, calibrated against a glass bottom dish of the same kind, and placed at 15 mm above each cell navigated by 405 nm epifluorescence and phase-contrast (Nikon Eclipse Ti-U with C-HGFI illuminator). One indentation per cell was performed by controlling piezo displacement for 1 mm indentation. The Young’s modulus was then derived along the loading curve from 0 to 1 mm using Hertzian contact model with Poisson’s ratio n = 0.5 in the Optics11 DataViewer software.

### Immunofluorescence and imaging

Cells were fixed with 3.7% paraformaldehyde (PFA) at room temperature for 10 minutes and then permeabilized and blocked with 0.1% Triton X-100 and 1% bovine serum albumin (BSA) in PBS at room temperature for 1 hour. For immunostaining, primary and secondary antibodies were prepared in a 1% BSA solution containing 0.1% Tween-20 in PBS (PBST). Cells were incubated overnight at 4 ^°^ C with the primary antibody against NFATC2 (GeneTex, 1:200, GTX127932). Following three washes with 0.1% PBST, cells were incubated for 4 hours at room temperature with the secondary antibody solution (biotium, Goat Anti-Rabbit IgG, 1:1000) and Hoechst 33342 (1:1000). Widefield fluorescence microscopy was performed using the ZEISS Celldiscoverer 7 microscope with a Plan-APOCHROMAT 20×/0.7 NA autocorrective objective.

### RNA sequencing

Total RNA was isolated and purified from cell pellets collected after mechanical dosing using the Arcturus PicoPure RNA Isolation Kit (Applied Biosystems). RNA concentration was quantified using a NanoDrop 8000 Spectrophotometer (Thermo Fisher Scientific). Bulk RNA sequencing was performed by BGI Transcriptome Sequencing. Differential gene expression analysis was conducted using DESeq2, and Gene Ontology (GO) enrichment analysis was performed using the enrichGO function from the clusterProfiler package in R. Differentially expressed genes were identified with the threshold of padj < 0.05 and |log2FoldChange| > log2(1.5). KEGG pathway enrichment analysis was performed using the enrichKEGG function from the clusterProfiler package, with padj < 0.05.

### NFATC2 knockdown

Cells were transfected with NFATC2 siRNA (30 pmol per sample, Silencer Pre-Designed siRNA, Thermo Fisher) or scrambled siRNA (30 pmol per sample, Negative Control #4 siRNA, Thermo Fisher) every two days using Lipofectamine RNAiMAX Reagent (7.5 µL per samples, Invitrogen).

Knockdown efficiency was assessed by SDS-PAGE and western blotting. Cell pellets were harvested after transfection, washed with PBS, and lysed in RIPA buffer supplemented with protease and phosphatase inhibitors (Thermo Fisher). Lysates were vortexed and centrifuged at 12,000 × g for 20 minutes at 4 ^°^ C. Supernatants were collected and protein concentrations were determined using a bicinchoninic acid (BCA) assay (Thermo Fisher). Equal amounts of protein were mixed with Laemmli sample buffer, denatured at 95^°^ C for 5 minutes, and separated on an 8% SDS-PAGE gel. Proteins were transferred to a PVDF membrane (GE Healthcare Life Sciences) using a wet transfer system (15 V, overnight). Membranes were blocked with 5% BSA in TBST for 1 hour, followed by overnight incubation at 4^°^ C with the primary antibody against NFATC2 (GeneTex, GTX127932) and GAPDH (ThermoFisher, PA1-987) diluted 1:1000 in 5% BSA in TBST. After washing, membranes were incubated with HRP-conjugated secondary antibodies (Invitrogen) for 1 hour at room temperature. Protein bands were detected using enhanced chemiluminescence (ECL, Bio-Rad) and imaged with a Bio-Rad ChemiDoc Touch Imaging System.

### Statistical Analysis

All data were analyzed in GraphPad Prism 9.5 and reported as mean ± 95% confidence interval. One-way ANOVA with Dunnett’s post-hoc test or Kruskal-Wallis with Dunn’s post-hoc test were used when appropriate. Two-way ANOVA with Šídák post-hoc test was used for the cell spreading assay. P-values <0.05 were labelled significant (^*^).

## Key Resources Table

| REAGENT OR RESOURCE | SOURCE | IDENTIFIER |
| --- | --- | --- |
| Antibodies |  |  |
| NFATC2 antibody | GeneTex | AB_2886213 |
| GAPDH antibody | ThermoFisher | AB_2107310 |
| Goat anti-rabbit IgG (H+L), CF488A-conjugated | Biotium | AB_10853132 |
| Goat anti-rabbit IgG (H+L), HRP | Invitrogen | AB_228341 |
| Bacterial and virus strains |  |  |
| Biological samples |  |  |
| Chemicals, peptides, and recombinant proteins |  |  |
| Collagen-1 | Gibco | A1048301 |
| Acrylamide/bisacrylamide | Bio-Rad | 1610140,<br>1610142 |
| Sulfo-SANPAH | ThermoFisher | 22589 |
| Critical commercial assays |  |  |
| Arcturus PicoPure RNA Isolation Kit | ThermoFisher | KIT0204 |
| BCA Protein Assay | ThermoFisher | 23225 |
| Deposited data |  |  |
| RNA-seq raw data | NCBI Sequence<br>Read Archive<br>(SRA) | PRJNA1258606 |
| Experimental models: Cell lines |  |  |
| HFF-1 | ATCC | CVCL_3285 |
| HT1080 | ATCC | CVCL_0317 |
| MDA-MB-231 | ATCC | CVCL_0062 |
| Experimental models: Organisms/strains |  |  |
| Oligonucleotides |  |  |
| NFATC2 siRNA | ThermoFisher | AM16708 |
| Negative control #4 siRNA | ThermoFisher | AM4641 |
| Recombinant DNA |  |  |
| LifeAct-TdTomato | Addgene | Addgene_64048 |
| Software and algorithms |  |  |
| GraphPad Prism 9.5 |  | SCR_002798 |
| DESeq2 |  | SCR_015687 |
| clusterProfiler |  | SCR_016884 |
| Ibidi Chemotaxis & Migration Tool V2.0 |  | SCR_017322 |
| Other |  |  |
| Zeiss Celldiscoverer 7 microscope | Zeiss |  |
| Optics11Life Chiaro nanoindenter | Optics11Life |  |
| Sony SH800S Cell sorter | Sony |  |
| Bio-Rad ChemiDoc Touch | Bio-Rad |  |

