## Supporting Information for "Mechanical Memory Primes Cells for Confined Migration"

**This file includes:**

Figs. S1 to S4

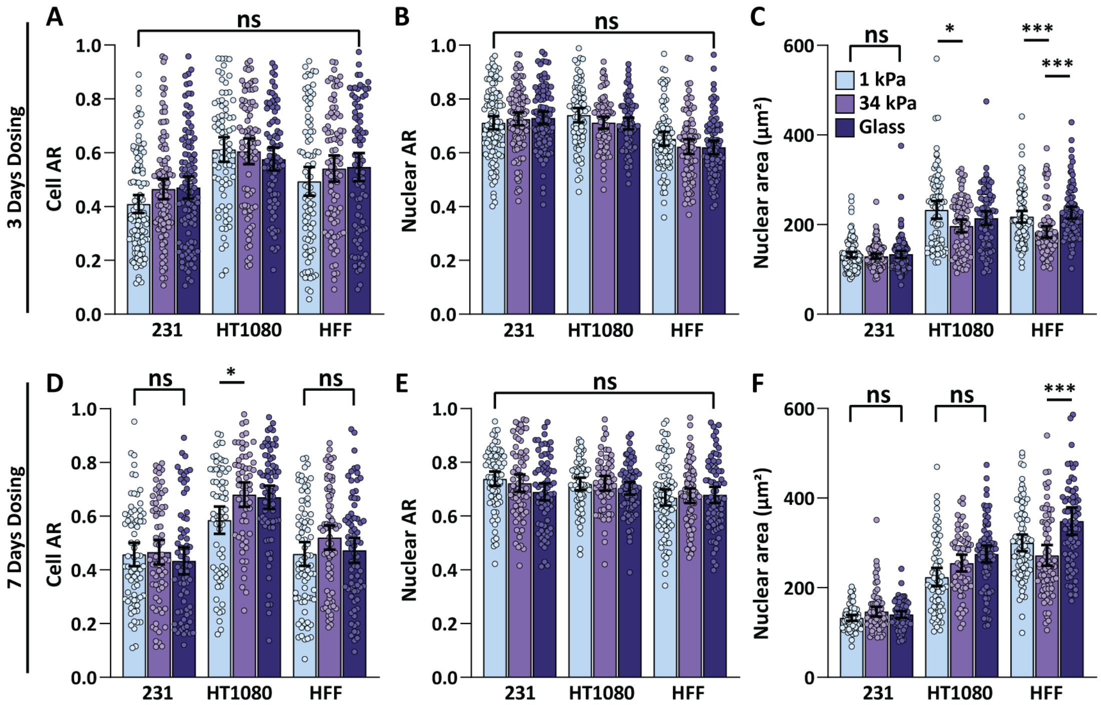


**Figure S1. Effects of Mechanical Dosing on Cell and Nuclear Morphology. (A to C)** Cell aspect ratio (AR), nuclear aspect ratio (AR), and nuclear area measured after 3-day mechanical dosing on hydrogels of varying stiffness. **(D to F)** Cell AR, nuclear AR, and nuclear area measured after 7-day mechanical dosing on hydrogels of varying stiffness. Results represented as mean ± 95% C.I., and ***P < 0.001, *P < 0.05 by Kruskal-Wallis and Dunn's post-hoc test.

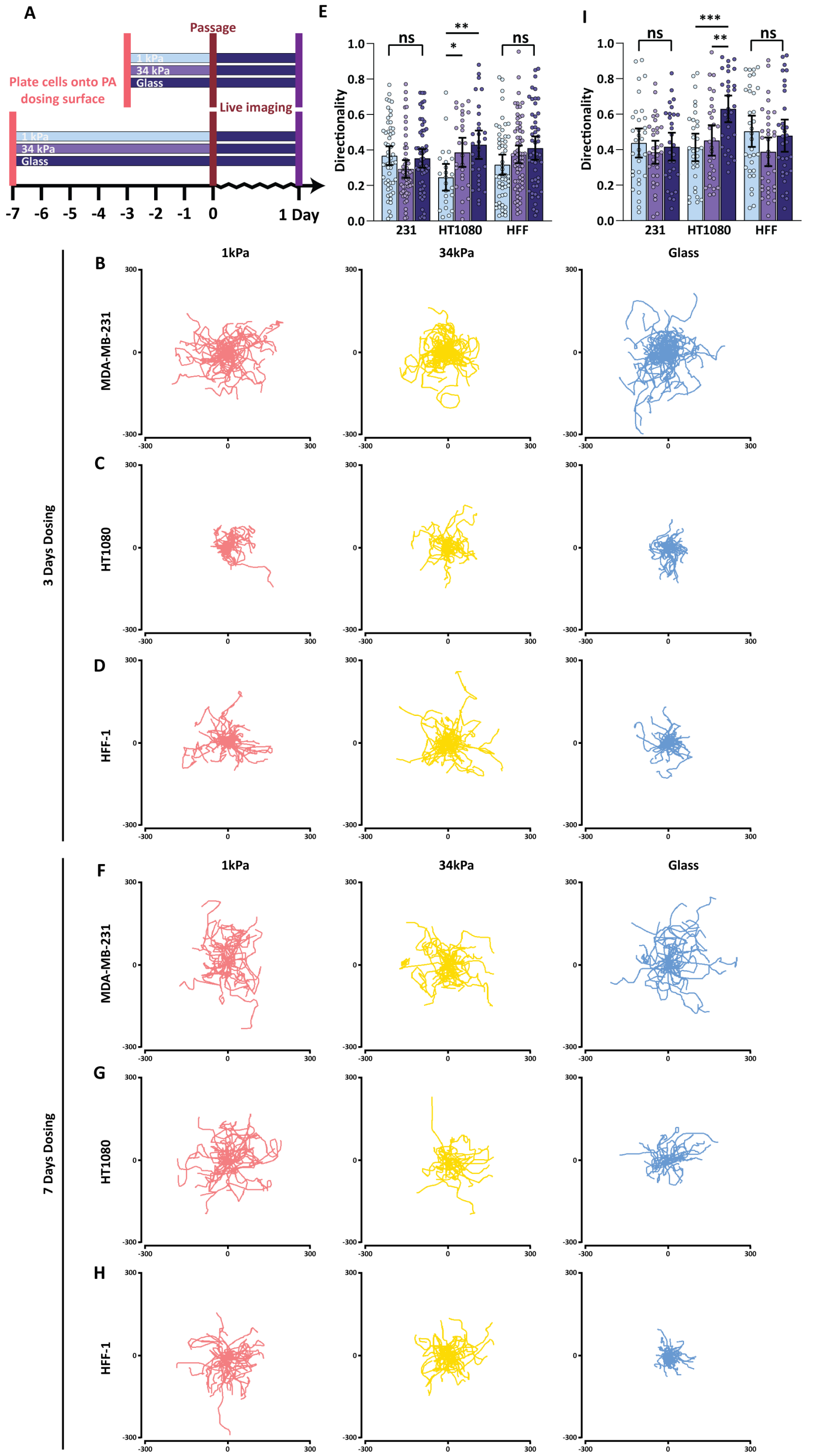


**Figure S2. Mechanical Dosing on Varying Stiffness Alters 2D cellular migration. (A)** MDA-MB-231, HT1080, and HFF-1 cell lines were cultured on PA hydrogels of different stiffness for 3 or 7 days prior to transfer onto glass surfaces for 2D migration tracking. **(B to E)** 2D migration directionality and migration trajectories of the three cell lines after 3-day mechanical dosing. **(F to I)** 2D migration trajectories and migration directionality of the three cell lines after 7-day mechanical dosing. Results represented as mean ± 95% C.I., and ***$P < 0.001$, **$P < 0.01$, *$P < 0.05$ by Kruskal-Wallis and Dunn's post-hoc test.

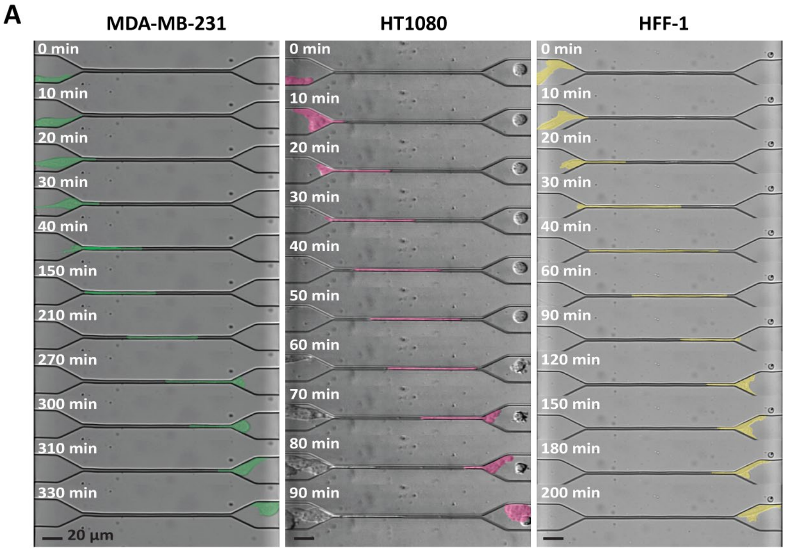


**Figure S3. Live-Imaged Microchannel Permeation Events for Confined Migration Speed Quantifications. (A)** Time-lapse images of MDA-MB-231, HT1080, and HFF-1 cells migrating through 3 µm-wide microchannels. Colored stamps indicate cell outlines.

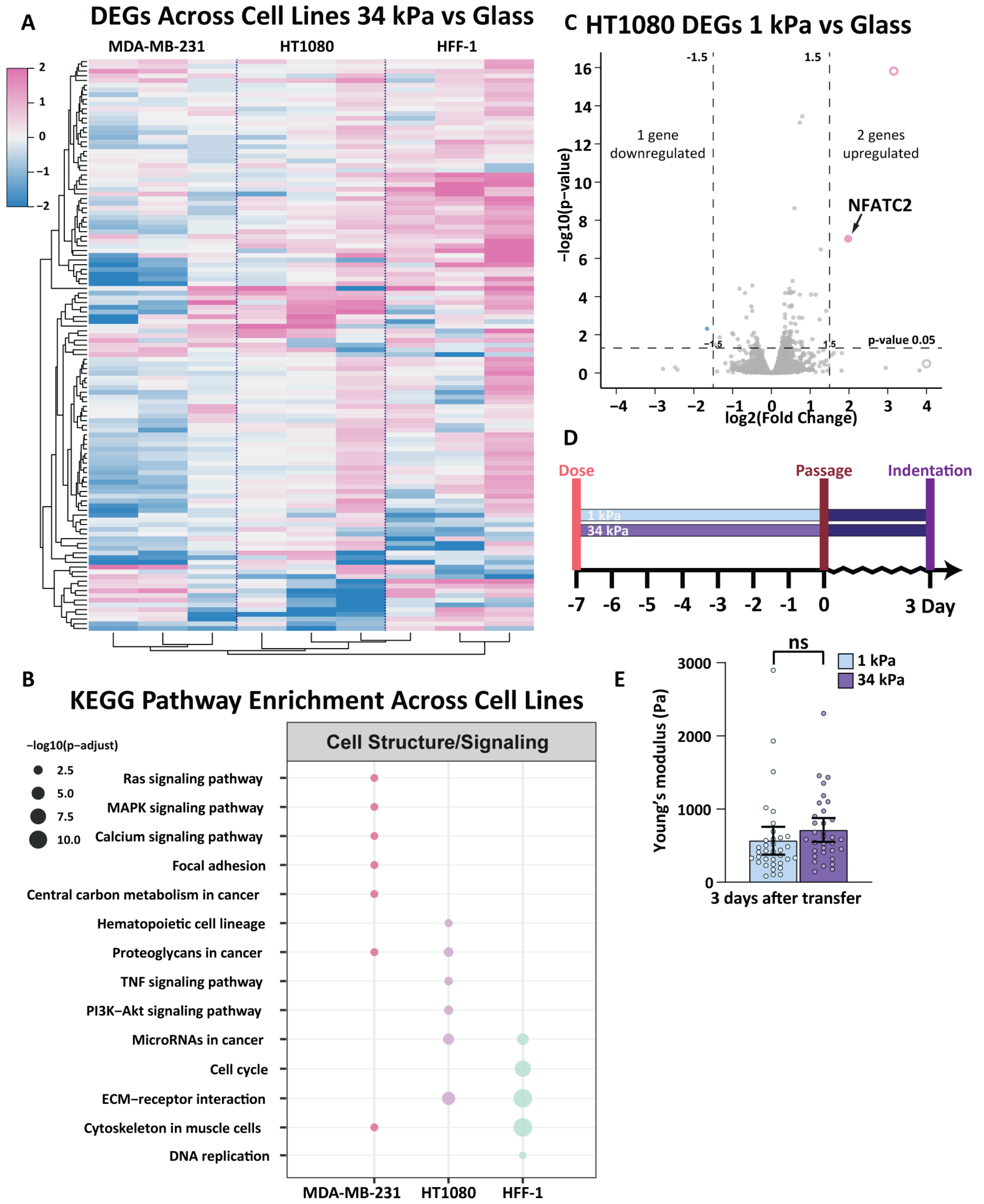


**Figure S4. Mechanical Dosing on Varying Stiffness Reprogrammed Cellular Transcriptional Activities and Mechanics. (A)** Heatmap showing the expression level of DEGs shared across cell lines resulting from 7-day mechanical dosing on 34 kPa hydrogel surfaces compared to glass. **(B)** Differentially enriched KEGG pathways from dosed MDA-MB-231, HT1080, and HFF-1 cells on 1 kPa hydrogel surface compared to glass. **(C)** Volcano plot showing differentially expressed genes between HT1080 cells dosed on 1kPa hydrogel surface and glass. Points cut off by axis were indicated by open circles. **(D)** Cells were dosed for 7 days followed by indentation after 3 days. **(E)** Cellular stiffness measured 3 days post-dosing. Results represented as mean ± 95% C.I..
